# CRISPR-Based Genome Editing in *Harmonia Axyridis*

**DOI:** 10.1101/2023.07.27.550814

**Authors:** Tamir Partosh, Michael Davidovitz, Noa Firer, Gur Pines

## Abstract

*Harmonia axyridis* (Pallas), commonly known as the Asian lady beetle, is a native insect species of Asia that has been intentionally introduced to various regions for biocontrol purposes. However, its widespread presence beyond its original release sites suggests a high degree of invasiveness. Previous studies have employed double-stranded RNA techniques to investigate the functional genomics of this species. In this study, we utilized the CRISPR-Cas9 approach to achieve precise genome editing in *H. axyridis*. Specifically, we targeted two distinct genes known to exhibit visible phenotypic effects. While the knockout of the *laccase2* gene resulted in an early-detectable phenotype but also in lethality, we successfully established a viable and genetically stable mutant colony by disrupting the *scarlet* gene. Our findings contribute to the expanding knowledge of genetic manipulation in *H. axyridis* and provide insights into its potential for future research and practical applications for biocontrol and invasive species management.

## 1. Introduction

*Harmonia axyridis*, also known as the Asian Lady beetle, is a highly polyphagous species with pray recorded to span several insect families, including Aphididae, Pseudococcidae, Aleyrodidae, Psyllidae, Coccinellidae, and other groups, with overall of more than 60 pray species, mostly aphids (van Driesche et al., 2018). As such, *H. axyridis* was extensively used for biocontrol purposes across the globe and is still being used in several locations. However, *H. axyridis* also displays high invasiveness properties, spreading beyond the original release sites (Kenis et al., 2020; Koch, 2003; Lombaert et al., 2010; Pines et al., 2022).

*H. axyridis* has four primary phenotypic forms; three include a black background with red or orange spots (*Forma conspicua, F. spectabilis, and F. axyridis*), whereas the fourth form (*F. succinea*) displays black spots over a red background (Michie et al., 2010). While the red and orange colors are nutritionally obtained and derived from their prey, the black color was shown to be self-synthesized melanin (Bezzerides et al., 2007).

The relative ease of CRISPR-Cas systems-based genome editing has made it possible to edit the genomes of organisms outside the traditional model organism playground. Indeed, newly edited organisms are being added to this ever-growing list, including insects. One of the main challenges of editing insect genomes is the delivery of the CRISPR-gRNA complex to the egg, which depends on a myriad of parameters, including the exact timing of microinjection, penetration technique, injection location, and post-injection conditions. Usually, the first targets for editing in insects are genes related to pigmentation, either throughout the body or in the eyes, which are easy to score and provide an estimate for the efficiency of the editing protocol (Reding et al., 2023).

In insects, the melanization pathway of the cuticle is conserved among species (Sugumaran and Barek, 2016). Various melanin molecules, including dopa, dopamine-melanin, N-acetyldopamine (NADA), and N-alanyldopamine (NBAD), are synthesized from a tyrosine precursor. *Laccase2* (*lac2*), a three-domain multi-copper oxidase, is the final enzyme in this pathway, as shown in the moth *Manduca sexta* and the beetle *Tribolium castaneum* (Arakane et al., 2009; Asano et al., 2019; Suderman et al., 2006). Regarding *H. axyridis*, several RNAi-based studies tested the functionality of aspartate decarboxylase, DOPA decarboxylase, tyrosine hydroxylase, and *laccase2*. These studies, performed on pupae and adults, showed that the *lac2* knockdown resulted in a significant loss of melanin production, leading to a distinct phenotype. (Chen et al., 2019; Niimi and Ando, 2021; Xiao et al., 2020; Zhang et al., 2020).

Dopa decarboxylase (DDC) knockout via CRISPR-Cas9 in *H. axyridis* was also recently reported, but it is unclear whether a stable mutant line was successfully established (Wu et al., 2022).

Changes in insect eye color are another common pigment-related phenotype used in functional genetics studies. Historically, such phenotypes played a major role in the genetic research of insects, such as the discovery of sex-linked inheritance in Drosophila melanogaster. (Morgan, 1910). Ommochrome precursors are transported into the insect eye via ATP-binding cassette transporters (ABC transporters) heterodimers. In *D. melanogaster*, these transporters are formed by a universal subunit, White, that heterodimerizes with either brown or *scarlet* (Ewart and Howells, 1998). RNAi functional tests established similar roles for White and *scarlet* in *T. castaneum*, but brown knockdown did not result in pigmentation differences (Grubbs et al., 2015). RNAi studies in *H. axyridis* established that the *white* and *scarlet* genes knockdown results in pigment-less eyes. However, bioinformatics analyses were unable to identify the *brown* gene (Tsuji et al., 2018).

Here, we used CRISPR-Cas9 to successfully target *lac2* and *scarlet* in *H. axyridis* and generate mutant strains. The *lac2* mutant allele is recessive lethal, but the phenotype was visible immediately after hatching, facilitating rapid editing efficiency estimations. Conversely, while the phenotype of *scarlet* knockout could only be identified in adults, homozygous mutants are viable and fertile, and a stable line bearing the phenotype was established.

## 2. Materials and Methods

### 2.1. Insect rearing

5-10 individuals of both sexes were grown in 90mm Petri dishes. A sugary food source in the form of a honey and paper towel mixture was always present on each plate. *Ephestia kuehniella* eggs or *Myzus persicae* were used as a protein-based food source. Each plate had 2-3 1.5ml plastic tubes to facilitate egg-laying.

For egg-laying synchronization, the protein-based food source was removed overnight and reintroduced in the morning. Egg collection was performed every hour by removing the 1.5ml tubes. This approach achieved up to six egg-laying cycles that started at around noon.

Post injection and hatching, individual larvae were reared in 60mm plates, with protein-based and sugary food sources present at all times. All insects were kept at 25 °C and 16 hours of light.

### 2.2. Primers and gRNA design

The *lac2* gRNA was designed based on the sequence obtained by Zhang et al. (Zhang et al., 2020): First, *Halac2* mRNA transcript (GenBank: MN650656.1) was BLAST-searched in the *H. axyridis* genome (icHarAxyr1.1). A single result aligned the coding sequence across several loci in chromosome 5 of the genome assembly (GenBank: OU611931.1).

The first exon sequence was extracted with an additional 500bp flanking the exon. PCR primers were designed to amplify an amplicon 486bp containing the exon (Table 1).

**Table 1.**
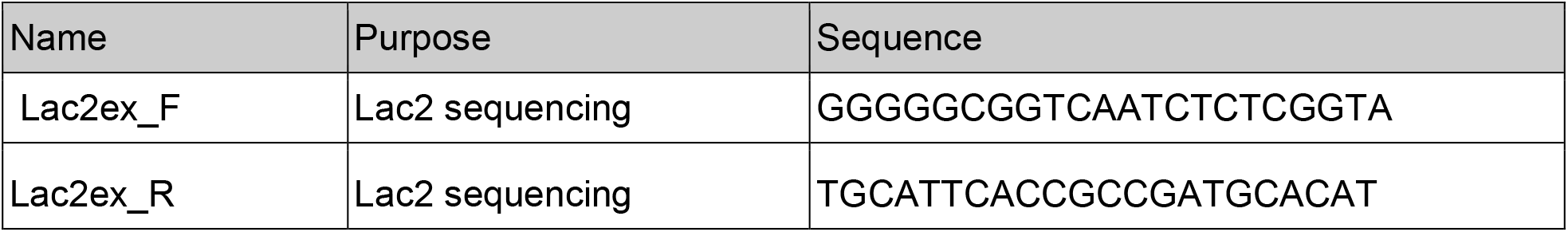
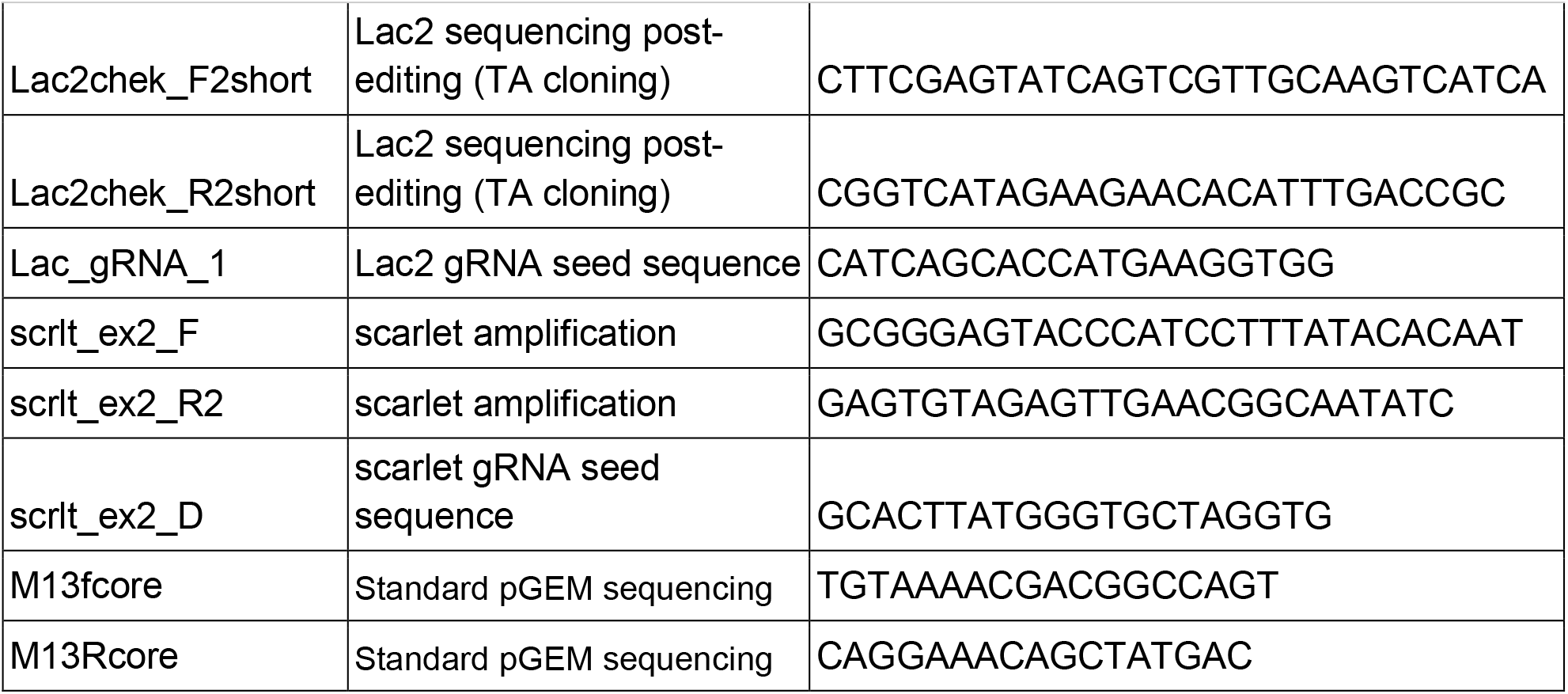
Primers used in this study.

Genomic DNA samples from eight *H. axyridis* adults were amplified, sequenced, and aligned to identify conserved regions vs. SNPs. Then, potential gRNA targets were identified using Benchling (benchling.com).

The list of target sequences was tested for off-target effect by performing another BLASTn run, and combined with the Benchling efficiency score, eight potential targets remained. From these, five were selected based on their location in the exon.

The five selected gRNAs were synthesized as Alt-R CRISPR-Cas9 crRNA by IDT, of which one was used for experiments (Lac_gRNA_1).

A similar process was done to design gRNA for the *scarlet* gene: The transcript sequence was obtained from Tsuji et al. (Tsuji et al., 2018) and was validated by sequencing several local individuals. Primers for exons of *scarlet* were designed, and amplicons of 635bp from three *H. axyridis* adults were amplified (with primers scrlt_ex2_R2 and scrlt_ex2_F) and were tested for SNPs. Three gRNA were designed, one of which was used for experimentations (scrlt_ex2_D). The primers and gRNA seed sequences used in this study are listed in Table 1.

### 2.3. Cas9/gRNA complexing

gRNA was synthesized by IDT. 1ul of a 100uM stock solution of both crRNA and tracer RNA were co-incubated in a thermocycler at 95C for 5 minutes and then set to cool at 25C for 5min. To form the ribonucleoprotein complex, 2ul of Cas9 (PnaBio, #CP01) 1000ng/ul were added and incubated with the gRNA at 25C for 5 minutes. 1ul of Injection buffer was added (5m MKCl and 0.1 mM phosphate buffer pH 6.8). The Cas9/gRNA mix was back-loaded to the capillary with Eppendorf Microloader pipette tips.

### 2.4. Microinjection

Borosilicate glass capillaries with filament (1×0.78mm, Warner Instruments,#G100TF-4) were used for microinjection, pulled to form injection needles in the “PUL-1000” pipette puller (World Precision Instruments). The pulling program was as follows: first two empty steps, then step 3 had heat-527, force-115, distance-1, and delay-0, followed by 4th step with heat-527, force-250, distance-1, and delay-10.

0-4h-old eggs were collected with a fine, moist brush and arranged on a 1*1 cm 3% agarose stair structure in straight rows along the grooves of the agarose structure, with each line having 10-15 eggs arranged in an anterior-posterior manner, with their lateral side fully exposed for injection. Injections were performed to the posterior third of the egg via the Injectman system (Eppendorf).

Post injection, the eggs were kept on the agarose structure for overnight incubation at 25 °C. The next day, each egg was placed separately in a 60mm new petri dish with 6-10 frozen *H. axyridis* eggs as a protein food source. The new hatchlings were fed daily until adulthood.

### 2.5. Mutant sequencing

Genomic DNA was obtained from exuviae samples collected through the beetles’ development through three molting phases and were kept at -20 °C for processing. DNA was extracted as previously described (Alon et al., 2023) with some adaptations:

Samples were placed in a 200ul PCR tube and were ground manually with 2ul of bio-grade water with the blunt end of a sterile plastic loop. Then, 20ul of saponin/PBS solution was added, homogenized with a motorized hand homogenizer, and incubated at 25 °C for 20 min. The tubes were then centrifuged at 18123G for 2 min, the supernatant was aspirated, and the pellet was washed with 20ul of PBS.

After washing, tubes were centrifuged again, the supernatant was aspirated, and 20ul of chelex 5% solution was added. The tube was vortexed for 10 sec, incubated at 100 °C for 10 min, and centrifuged again. The clear supernatant was measured in a spectrophotometer to estimate DNA quantity and was used in downstream applications.

The target cut sites were amplified by PCR using GoTaq G2 Green (Promega,# 9PIM782). Similar to Reding et al. (Reding and Pick, 2020) PCR product was A-tailed by incubation of the purified product with Taq polymerase, TA-cloned into the pGEM-T Easy vector (Promega), and Sanger-sequenced with the M13R primer (HyLabs, Israel).

## 2.6. Stabilization of scarlet mutant line and crossing scheme

out of 150 *H. axyridis* embryos treated with Cas9/scrlt_ex2_D, 14 reached adulthood. Two beetles with distinct white-eye phenotypes (Fig. S3) were copulated in a 90mm petri dish containing 1.5ml plastic tubes for egg-laying, a sugary food source, a folded wet paper towel, and a 2*2 cm bell pepper leaf containing 20+ aphids as a protein food source. Following egg hatching, Embryos were collected, and each G1 individual was moved to a separate 60cm petri dish and reared till adulthood. four mating pairs of G1 were copulated to form the G2 generation. Out of them, one G1 couple was substantially viable compared to the rest. That pair gave rise to a second generation (G2), in which both males and females had distinct white-eye phenotypes. Three mating pairs from G2 were formed to form G3 offsprings. G3 beetles were allowed to form mating pairs spontaneously in larger rearing groups. Additionally, two back-crossing pairs were copulated with G1 white-eye males and WT females. heterozygous offspring of the back-crossing pairs were coupled to get stable homozygous mutants. The crossing scheme is shown in Fig. 3C.

## 3. Results

### 3.1. lac2 knockout

Embryos were injected with the Cas9/gRNA RNP targeting the beginning of intron 1 (Fig. 1D). Larval hatching was monitored post-microinjection. Hatching started on the third-day post-injection and continued to day four. Edited larvae showed non-melanized patches in different sizes in all body parts, including the head, trunk, and limbs. The patchy pattern is likely due to the bi-allelic mosaicism as expected from injected individuals, in which mutation occurs only in some somatic cells. The phenotype was observable immediately after hatching and became clearer at later stages (Fig. 1A, B, C). Larval mortality correlated with phenotype intensity, and a severe phenotype was followed by early death up to several minutes following hatching (Fig. S1A, B). Genomic DNA was extracted from exuvia, and the *lac2* gene was sequenced to confirm successful genome editing at the desired locus: Sanger sequencing of individuals revealed that different alleles were present in each specimen, as expected, including disruption of the start codon and frameshifts (Fig. 1E).

**Figure 1.**
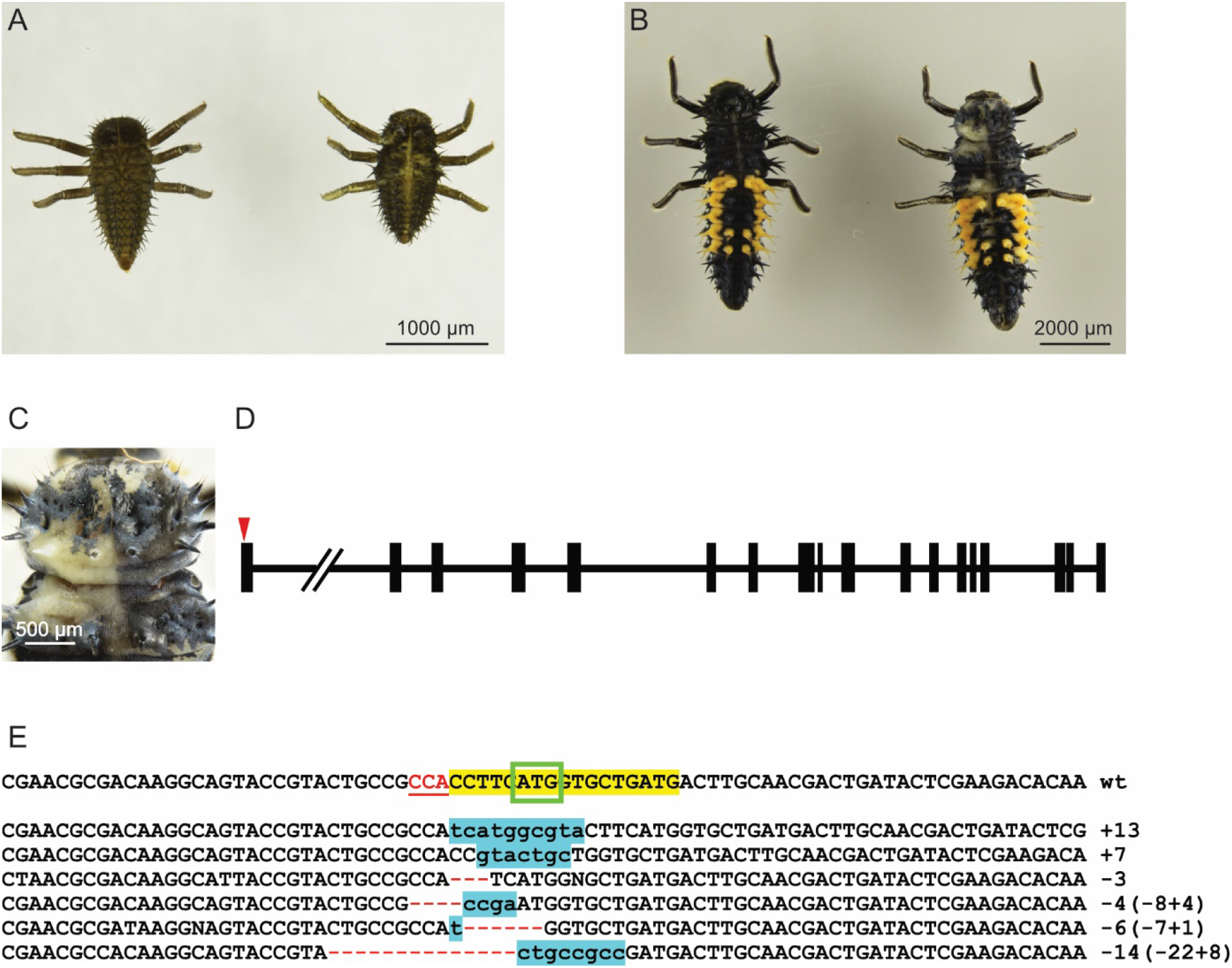
*lac2* knockout. (A), (B) Side-by-side comparison of wild-type (left) vs. G0 mutant (right) at 1st and 4th larval stages, respectively. (C) A zoom-in image on the edited larva from B, depicting the mosaic phenotype. (D) A schematic representation of the lac2 gene. Black vertical lines depict exons, and the horizontal lines are introns. The double slash indicates a long, not to scale intron. The red arrowhead indicates the gRNA targeting site. (E) Mutant sequences derived from G0 exuvia. The yellow highlight indicates the seed sequence, the red, underlined sequence represents the PAM sequence, and the green box marks the start codon. Light blue highlights indicate nucleotide insertions and red dashes represent deletions. The sum of the nucleotide changes is indicated on each mutant’s right.

Of all survivors bearing visible phenotypes, one male reached adulthood (Fig. S2A, B), and later sequencing of its gonads verified the presence of mutant alleles (Fig. S2C). This single adult was backcrossed to a wild-type female. Sequencing of G1 offspring genomic DNA confirmed that mutant alleles were transmitted, confirming successful germline editing (Fig. S2D). However, no G2 homozygous progeny displaying the expected phenotype could be found in the offspring of G1, probably due to lethality. Indeed, previous RNAi knockdown in *T. castaneum* showed structural cuticle abnormalities and premature death (Arakane et al., 2005).

### 3.2. scarlet knockout

To achieve our goal of establishing a stable gene-edited line, we next targeted the *scarlet* gene, whose knockout should result in a white-eye phenotype. gRNA sequences were designed according to the sequence reported by Tsuji et al. (Tsuji et al., 2018), and targeted exon 14 (Fig. 2C). Embryo injection was conducted similarly to the *lac2* injections. Surviving injected larvae showed no apparent morphological difference from buffer-injected control individuals. Two adults, a male and a female, emerged with complete or partial white eyes (Fig. S3), and were crossed with each other. An example of white-eyed *H. axyridis* vs. wild-type is shown in Figure 2B. Exuvia-derived genomic DNA showed the expected short insertions and deletions that would be expected from non-homologous end-joining repair of double-strand breaks (Fig. 2D).

**Figure 2.**
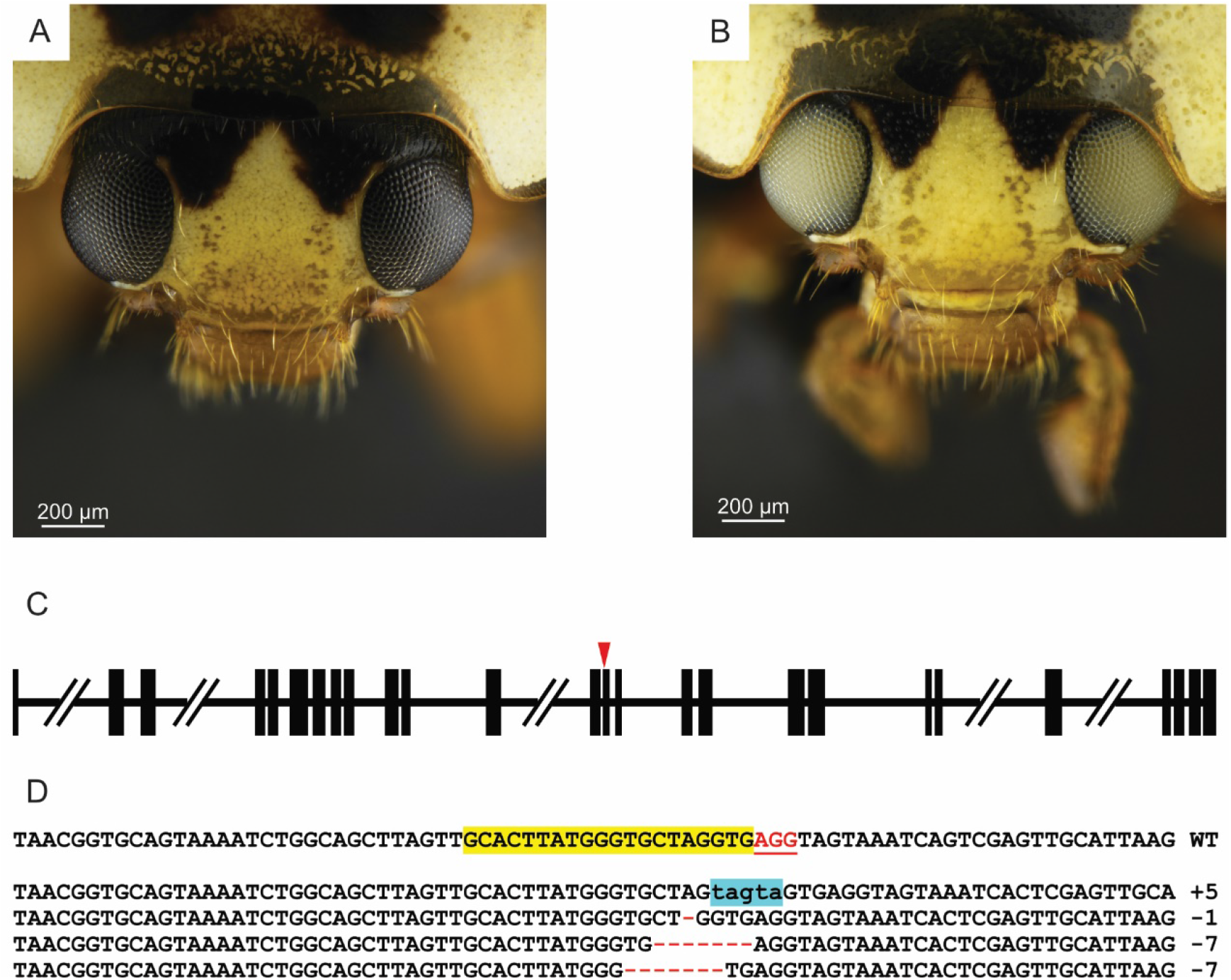
*scarlet* gene knockout. (A), (B) A close-up image of a wild type and mutant *H. axyridis* adult, respectively. C. a schematic representation of the *scarlet* gene. Black vertical lines depict exons, and the horizontal lines are introns. The double slashes indicate long, not to scale introns. The red arrowhead indicates the gRNA targeting site. (D) Mutant sequences derived from G0 exuvia. The yellow highlight indicates the spacer sequence, and the red, underlined sequence represents the PAM sequence. Blue highlights indicate nucleotide insertions, whereas red dashes represent deletions. The sum of the nucleotide changes is indicated on each mutant’s right.

**Figure 3.**
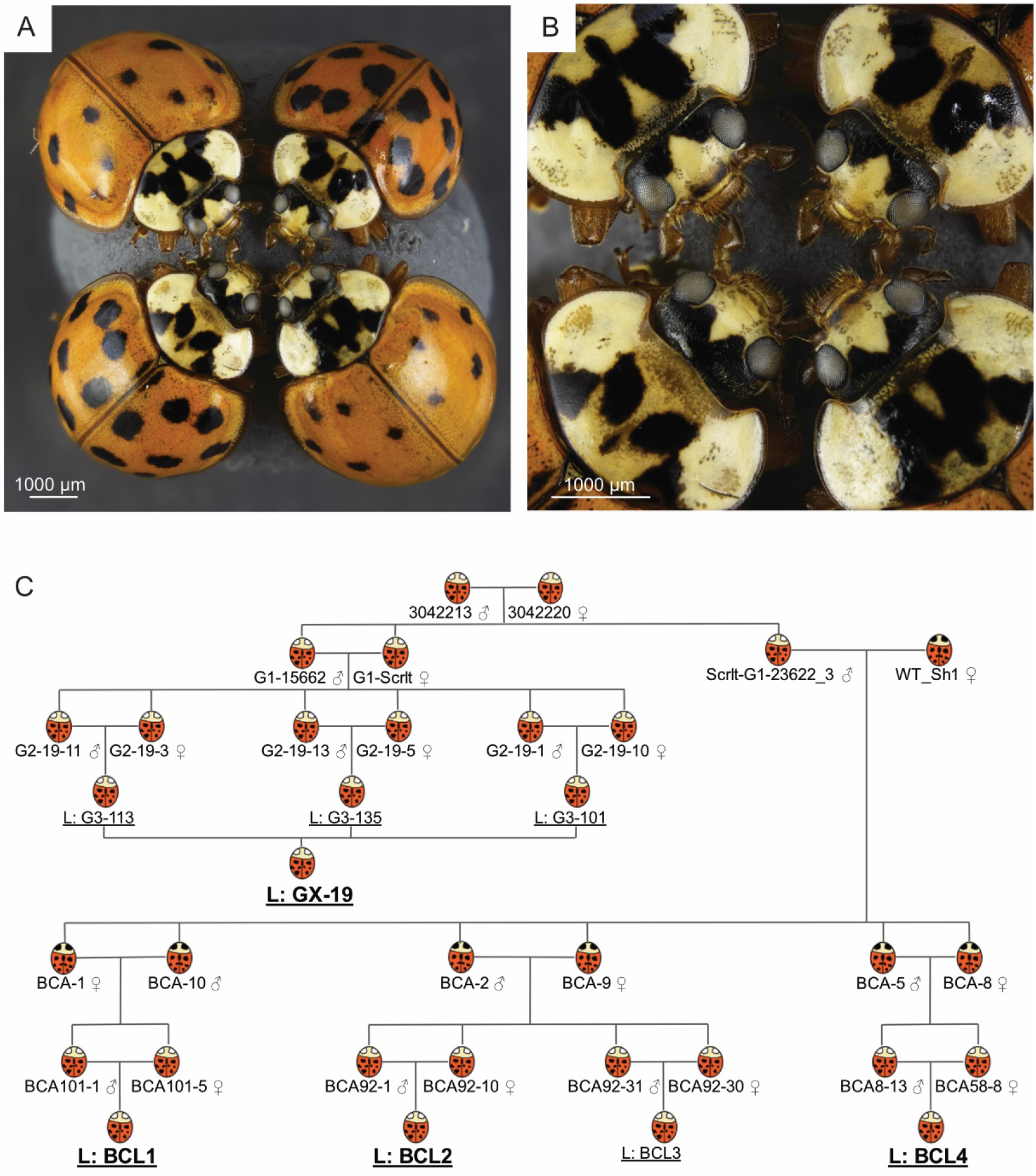
Establishing a stable white-eyed mutant line. (A) Four representative specimens from the GX-19 line. (B) Zoom-in of A, highlighting the consistent white-eye phenotype. (C) A schematic plot representing the crossing scheme used to establish the four stable lines highlighted in bold. BCL, Back Cross Line. Black and white eyes represent wild-type and mutant phenotypes, respectively.

All subsequent G1 offspring were white-eyed, suggesting successful germline editing. A G1 couple gave rise to six fertile and viable G2 parents, their G3 offspring forming the GX19 line (Fig. 3A, B, C). One G1 white-eyed male was coupled with a black-eyed WT female, giving rise to fertile and viable heterozygous black-eyed couples. G3crossing gave rise to a mixed-phenotype population, out of which only the white-eyed individuals were selected to form three mating pairs to establish the “Back-Cross” line (BCL) (Fig. 3C). This backcrossing established that the mutation is recessive since roughly 25% of the progeny had white eyes (Castle, 1903). The white-eyed colony has been maintained for over five generations with a stable phenotype in all subsequent offspring (Fig. 3A, B).

## 4. Discussion

Genome editing efforts currently exit the realm of model organisms and expand to previously considered complex and laborious organisms. Genome editing of *H. axyridis* was previously shown using the transcription activator-like effector nuclease (TALENs) approach to knock out an inserted transgene (Hatakeyama et al., 2016), and recently CRISPR/Cas9 method was used to knock out the dopa-decarboxylase (DDC) gene (Wu et al., 2022). However, it was unclear from that work whether or not a stable mutant line was established. Here we demonstrate the successful genome editing of the harlequin Asian ladybug *H. axyridis* utilizing the CRISPR/Cas9 system and the establishment of a stable mutant line. We edited two phenotypic genes to simplify screening and efficiency evaluations. However, while *lac2*, similarly to *ddc*, was shown to have a role in the melanin metabolic pathway, this gene also has a pivotal role in cuticle sclerotization, promoting protein-protein and protein-chitin cross-linkages (Asano et al., 2019).

Indeed, several insects treated with *lac2* dsRNA exhibit molting defects to the point of lethality in a dose-dependent manner, including *Tribolium castaneum, Nilaparvata lugens*, and *Riptortus pedestris* (Arakane et al., 2005; Futahashi et al., 2011; Ye et al., 2015). This central aspect of *lac2* may explain the high lethality we observed in the edited insects relative to controls (Fig. S1), and may also explain our unsuccessful efforts to generate mutant homozygotes. We, therefore, selected *scarlet* and successfully induced specific edits with a visible phenotype allowing us to establish a viable colony of beetles with distinguished white-eyed phenotype targeting the *scarlet* locus. The advantage of knocking out *lac2* is that a clear phenotype is readily available to assess the editing efficiency rapidly, as the phenotype is seen immediately after hatching. On the other hand, the *scarlet* knockout, through the establishment of a genomically edited line, exemplifies the potential for maintaining an altered trait across generations. These successful examples of Cas9-based genomic edits in *H. axyridis* pave the way for additional functional genomics studies of this ecologically-and agriculturally-important insect. Our editing protocol may be used to generate other edits in *H. axyridis* for functional genomics studies, agricultural biocontrol purposes, and the management of invasive species. Moreover, the *H. axyridis* white-eye mutations may serve as biomarkers for several applications, including screens for other mutations or, with an added fluorescent marker, play a role in pest management programs, as suggested before for fruit flies and other beetles (Berghammer et al., 1999; Kane et al., 2017).

## Supplementary figures

**Figure S1.**
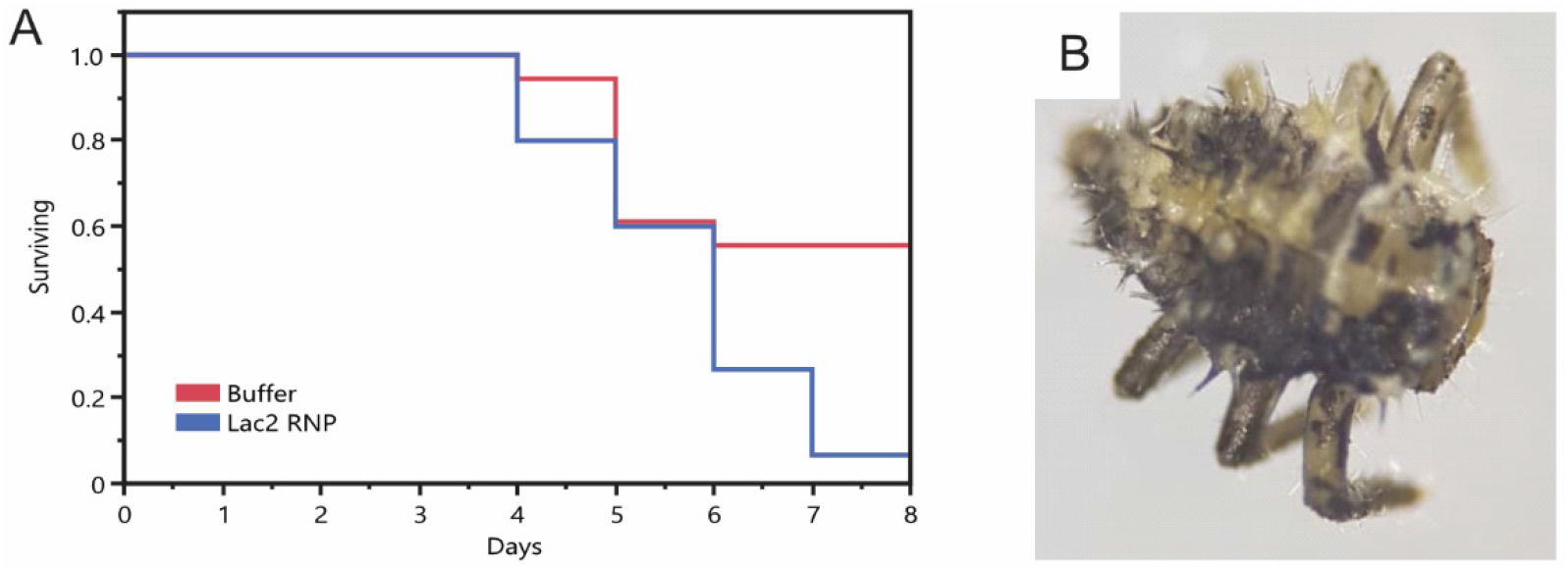
*lac2* knockout G0 adult. A single adult of the *lac2* RNP survived with a mild phenotype on the elytra. This specimen was edited in its gonads, and backcrossing resulted in wild-type and heterozygous progeny only, suggesting the *lac2* knockout homozygosity is lethal.

**Figure S2.**
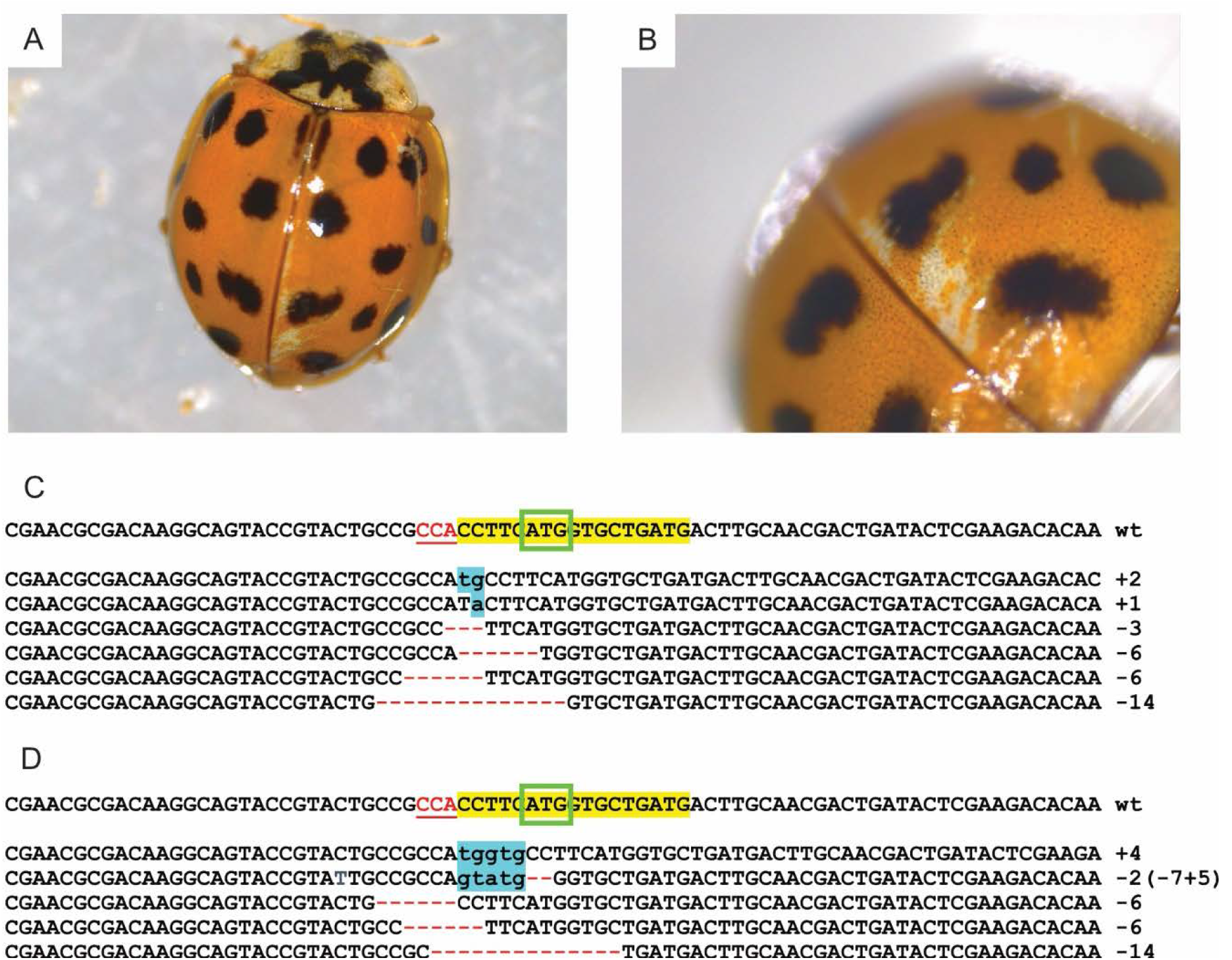
*lac2* adult mutant. (A) A single edited insect survived to adulthood with some phenotypic white patches on its elytra. (B) Zoom in on the white patch. (C) After the edited adult’s death, the gonads were surgically removed. TA cloning of the *lac2* amplicon confirmed germline edits. (D) Sequencing results of the G1 progeny of a cross between the edited adult male and a wild-type female.

**Figure S3.**
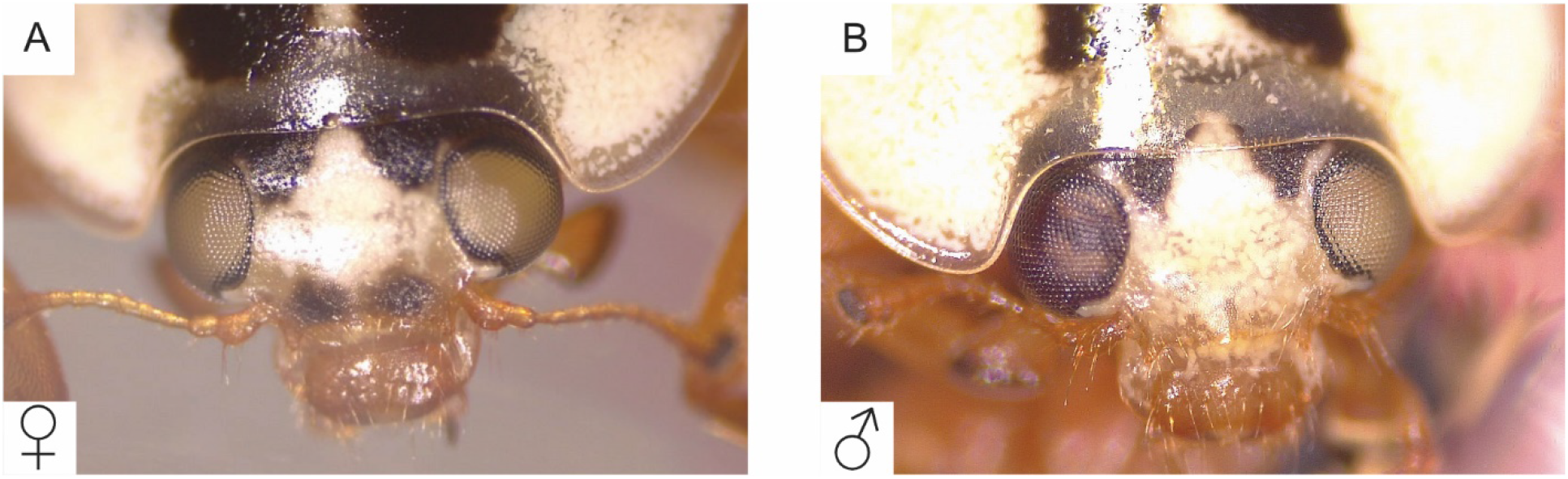
The G0 *scarlet* stable line founders. Two adults displayed a degree of white-eye phenotype, suggesting high editing efficiency. While the female’s eyes were completely white (A), the male displayed a classical mosaic pattern in one eye, while the other was almost entirely white (B).

## Acknowledgments

We thank Roy Kaspi for introducing us to this fascinating insect.

